# PROMO: An interactive tool for analyzing clinically-labeled multi-omic cancer datasets

**DOI:** 10.1101/629584

**Authors:** Dvir Netanely, Neta Stern, Itay Laufer, Ron Shamir

## Abstract

**Background:** Analysis of large genomic datasets along with their accompanying clinical information has shown great promise in cancer research over the last decade. Such datasets typically include thousands of samples, each measured by one or several high-throughput technologies (‘omics’) and annotated with extensive clinical information. While instrumental for fulfilling the promise of personalized medicine, the analysis and visualization of such large datasets is challenging and necessitates programming skills and familiarity with a large array of software tools to be used for the various steps of the analysis.

**Results:** We developed PROMO (Profiler of Multi-Omic data), a friendly, fully interactive stand-alone software for analyzing large genomic cancer datasets together with their associated clinical information. The tool provides an array of built-in methods and algorithms for importing, preprocessing, visualizing, clustering, clinical label enrichment testing and survival analysis that can be performed on a single or multi-omic dataset. The tool can be used for quick exploration and for stratification of tumor samples taken from patients into clinically significant molecular subtypes. Identification of prognostic biomarkers and generation of simple subtype classifiers are additional important features. We review PROMO’s main features and demonstrate its analysis capabilities on a breast cancer cohort from TCGA.

**Conclusions:** PROMO provides a single integrated solution for swiftly performing a complete analysis of cancer genomic data for subtype discovery and biomarker identification without writing a single line of code, and can, therefore, make the analysis of these data much easier for cancer biologists and biomedical researchers. PROMO is freely available for download at http://acgt.cs.tau.ac.il/promo/.

## Background

In recent years, a growing number of high-throughput genomic technologies have become available for biomedical research and are jointly providing high-resolution genomic data that fuel the revolution of personalized medicine [1][2]. These technologies (collectively named omics) allow the simultaneous quantification of a large number of features at various biological levels. The features include gene expression (mRNA and miRNA abundance levels measured by microarrays or RNA-Seq), protein expression (measured by mass spectroscopy or reverse phase protein arrays), DNA methylation (methylation arrays), copy number variation (SNP arrays), and others [3][4]. The technologies vary broadly in the number of features they measure as well as in the distribution of measured values [5]. However, they can typically be summarized as a numeric matrix where columns represent samples and rows represent biological features (often correlating to genes). Bioinformatic analysis of such genomic matrices has been extensively used for identifying biologically distinct sample groups, and for revealing groups of correlated biological features [6][7].

The number of tumor samples and measured features that are included in a typical cancer genomic dataset have grown dramatically in the last few years, owing to increasing resolution and reduced costs of array and sequencing technologies. Modern repositories comprise thousands of patient samples and many thousands of features. Investigation of such large datasets is computationally challenging as it requires robust software tools for supporting the analysis of both samples and features in high dimensional data [8].

In addition to genomic data, modern cancer datasets can include extensive medical information (labels) describing each sample, such as clinical properties or assignment to a predefined phenotype. These clinical labels make it possible to fuse genomic and clinical data in various ways in order to discover new insights based on feature-phenotype associations. Common clinical labels in cancer datasets include disease subtypes, pathological stages, survival and recurrence follow-up information, as well as response to treatment. Identification of genomic features that are correlated with significant clinical parameters (biomarkers) is expected to play a significant role in the field of personalized medicine, by which the status of multiple biomarkers may improve subtype diagnosis and guide therapeutic decisions [9][10].

The Cancer Genome Atlas (TCGA) is an example of a revolutionary multi-label multi-omic genomic database [11]. It includes more than 11,000 samples from 33 types of cancer, where each sample was measured using multiple omic technologies and was described by dozens of clinical labels [12]. Many studies have already analyzed TCGA data, improving the subtyping of cancers and shedding light on the biological mechanisms underlying the development of various cancer types [13][14][15]. Such analyses are typically time-consuming, computationally challenging, and entail team effort, as they require applying a diverse array of methods, statistical tools, and algorithms, and often also require writing extensive computer code to perform and interweave the various steps of the analysis [16]. Hence, to effectively extract clinically meaningful insights from such multi-omic multi-label databases, specialized agile integrative tools are required.

To address this challenge, we developed **PROMO** (**PRO**filer of **M**ulti **O**mic data), a fully interactive software suite capable of quickly importing, preprocessing, visualizing, analyzing and reporting the results on cancer datasets in a seamless fashion, without writing a single line of computer code. PROMO includes an extensive array of bioinformatic methods for performing major common analysis types including exploration, visualization, identification of clinically significant disease subtypes, revealing co-regulated feature groups, biomarker discovery, simple classification and integrative multi-omic analysis. Table 1 presents an overview of the fundamental analysis types available in PROMO.

**Table 1.**
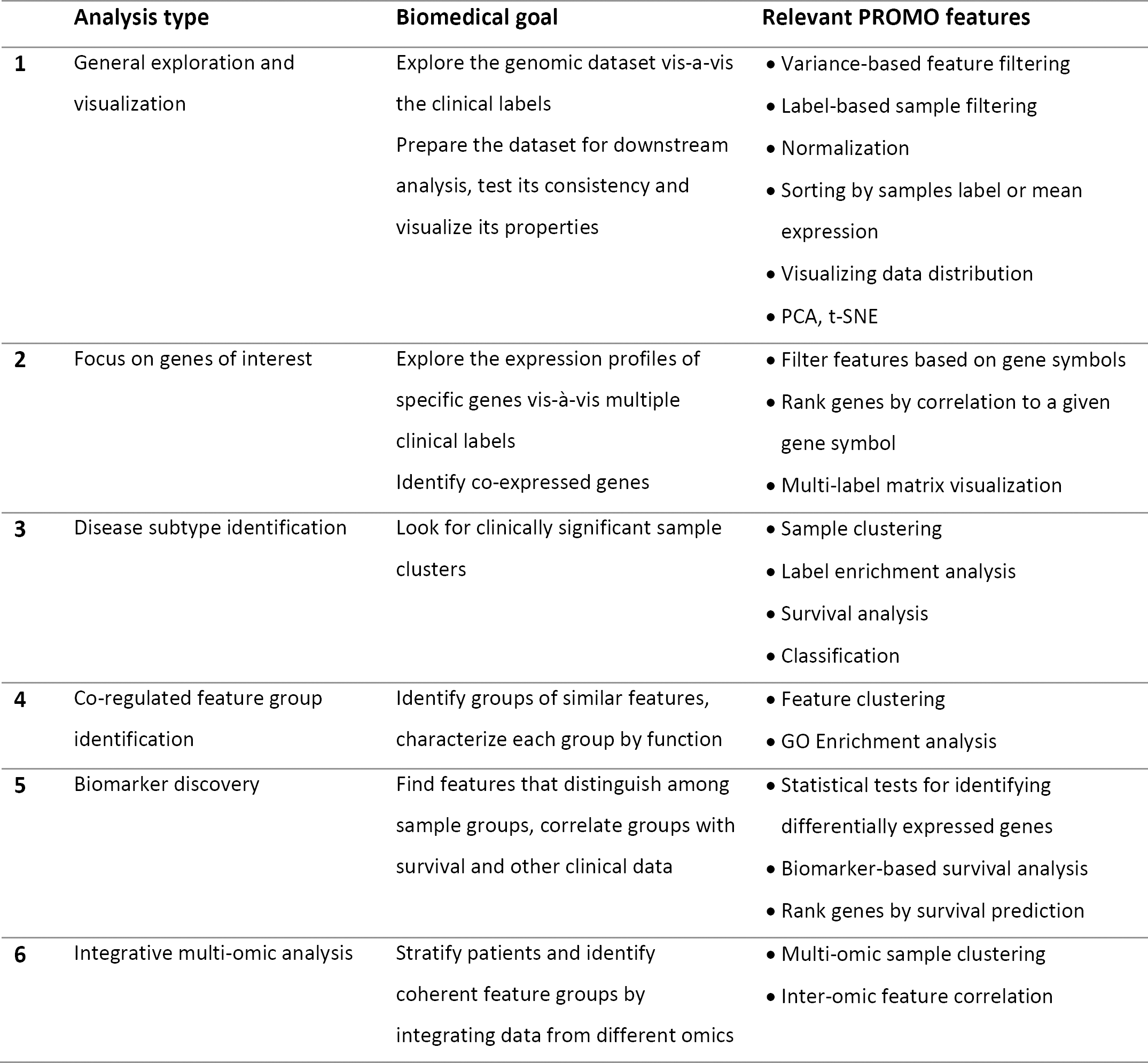
PROMO’s main analysis types

|  | **Analysis type** | **Biomedical goal** | **Relevant PROMO features** |
| --- | --- | --- | --- |
| **1** | General exploration and visualization | Explore the genomic dataset vis-a-vis the clinical labels<br>Prepare the dataset for downstream analysis, test its consistency and visualize its properties | • Variance-based feature filtering<br>• Label-based sample filtering<br>• Normalization<br>• Sorting by samples label or mean expression<br>• Visualizing data distribution<br>• PCA, t-SNE |
| **2** | Focus on genes of interest | Explore the expression profiles of specific genes vis-à-vis multiple clinical labels<br>Identify co-expressed genes | • Filter features based on gene symbols<br>• Rank genes by correlation to a given gene symbol<br>• Multi-label matrix visualization |
| **3** | Disease subtype identification | Look for clinically significant sample clusters | • Sample clustering<br>• Label enrichment analysis<br>• Survival analysis<br>• Classification |
| **4** | Co-regulated feature group identification | Identify groups of similar features, characterize each group by function | • Feature clustering<br>• GO Enrichment analysis |
| **5** | Biomarker discovery | Find features that distinguish among sample groups, correlate groups with survival and other clinical data | • Statistical tests for identifying differentially expressed genes<br>• Biomarker-based survival analysis<br>• Rank genes by survival prediction |
| **6** | Integrative multi-omic analysis | Stratify patients and identify coherent feature groups by integrating data from different omics | • Multi-omic sample clustering<br>• Inter-omic feature correlation |

An early version of PROMO was developed as part of a study where we identified distinct prognostic subgroups in Luminal-A breast tumors based on expression and methylation data [17]. The analysis workflow in that project provides an example of the key steps in a typical application of PROMO (Figure 1): Data are imported, filtered and preprocessed. Tumor samples are clustered into groups that are then assessed for clinical significance using survival analysis and statistical tests on the clinical labels. Clustering of the genes followed by gene enrichment analysis, associates sample clusters with active gene functions. The analysis is summarized visually in a genomic matrix clearly showing the identified sample clusters and their association to important clinical labels (Figure 1, step 4), in addition to downstream analysis methods (Figure 1, steps 5-7).

**Figure 1.**
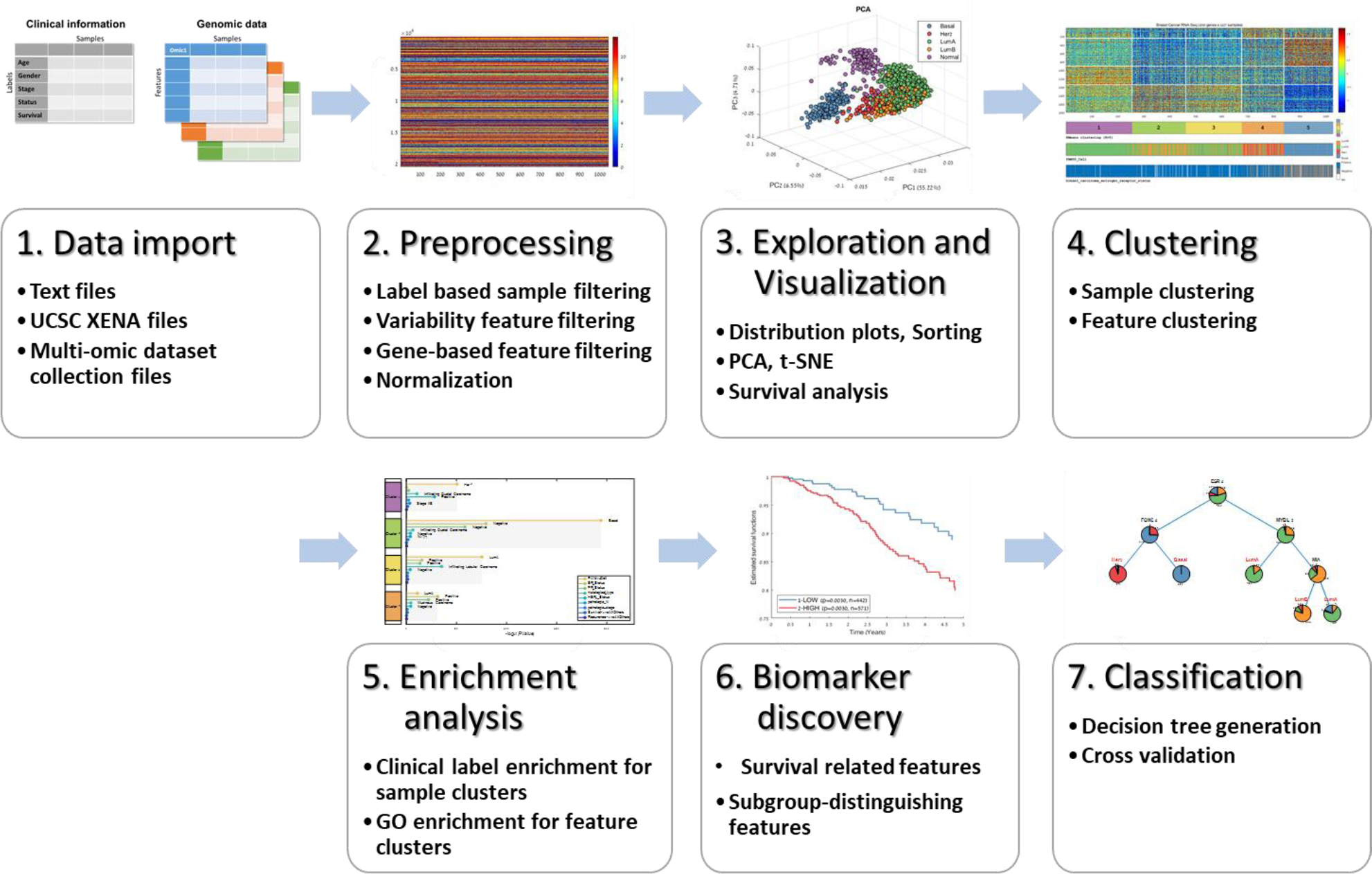
PROMO’s subtype discovery workflow – From data import to subtype classifier. This figure outlines the complete workflow by which PROMO can be used for identifying and characterizing clinically distinct cancer subtypes: (1) Importing genomic data together with clinical information in one of several available formats. (2) Preprocessing the data and preparing it for downstream analysis. (3) Verifying the integrity of the data, characterizing its distribution and exploring dataset properties with respect to available clinical label. (4) Employing clustering algorithms partition both samples and features (genes) into groups. (5) Applying enrichment tests to identify clinically significant sample subtypes and groups of co-regulated genes. and to characterize their function. (6) Statistical tests identify features that distinguish between different sample subtypes as well as survival related features. (7) Decision tree classifier can be generated for formulating a set of rules by which a new sample can be classified.

In this paper, we describe PROMO’s main features and demonstrate its use in a study of a breast cancer cohort [14].

## Implementation

PROMO is a standalone Windows application that can support huge datasets and has a fast fully interactive graphical user interface. PROMO was written in MATLAB, and it runs over the freely available Matlab runtime environment, taking advantage of its strong computational engine and editable graphical outputs. PROMO is freely available for download at http://acgt.cs.tau.ac.il/promo/.

PROMO’s main screen (Figure 2A) includes several key graphic elements: A large heatmap representing the currently analyzed genomic matrix is located at the center of the screen (heatmap colors correspond to the matrix values as indicated by the color scale on the right). Beneath the heatmap, a color-bar displays the currently selected sample labels. The same sample label colors will consistently be used by PROMO in all displays. The user can scroll down the list of clinical labels and explore their distribution over the samples. The panel on the left provides access to common commands and parameters. A text log that documents the analysis steps appears at the bottom of the screen. Figures 2B-F show the various panels that can be directly opened from the tab menu on the left of the screen, providing quick access to PROMO’s most useful features.

**Figure 2:**
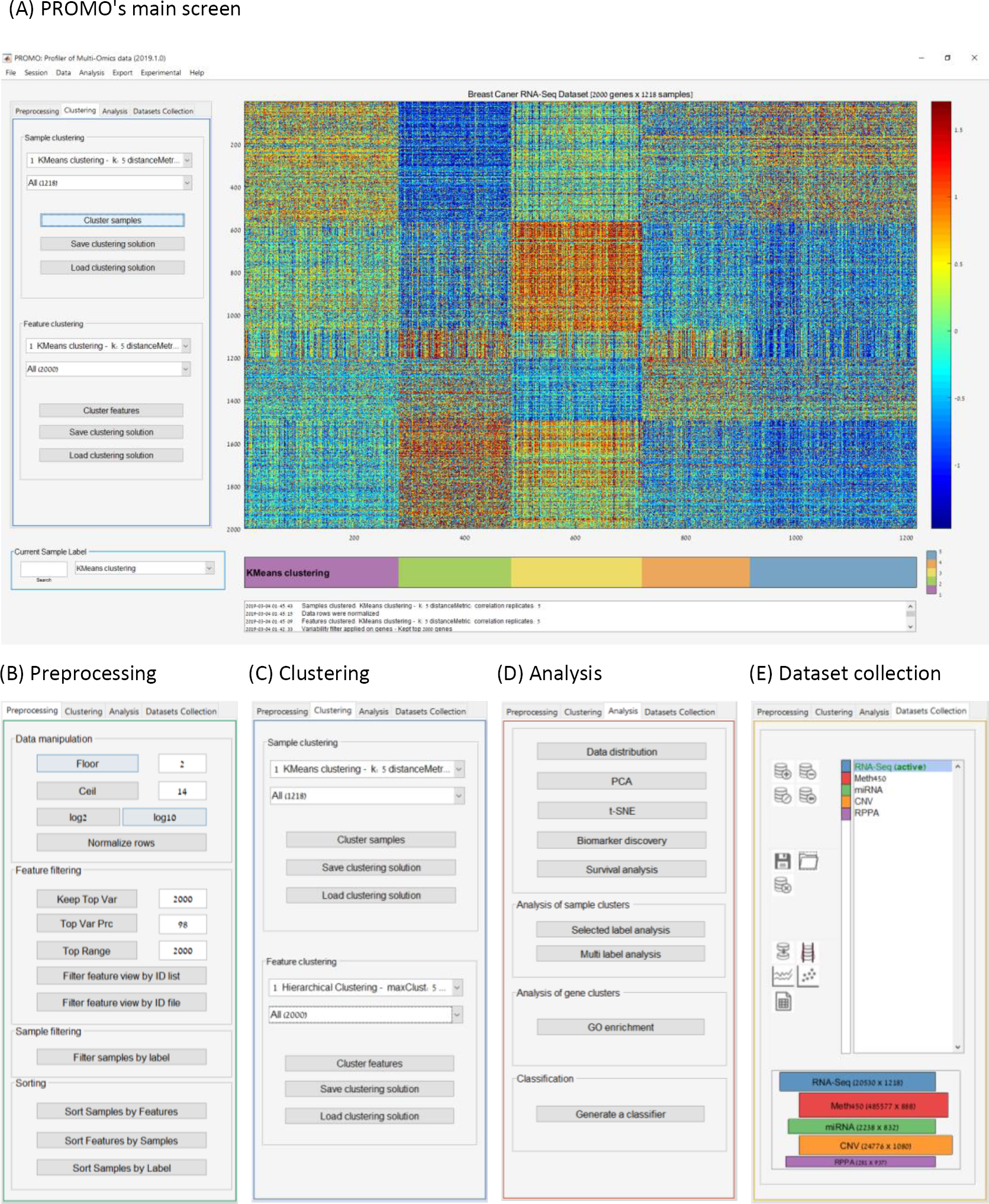
PROMO’s graphical user interface. (A) PROMO’s main screen. The genomic matrix is in the center with columns corresponding to samples and rows to features. Colors represent feature values according to the scale on the right. The colorful label bar beneath the matrix displays the currently selected sample label. Analysis steps are documented in the textbox on the bottom of the screen. Key commands are available on the tabbed panels on the left of the screen. (B) The *Preprocessing* panel allows filtering, normalization, and sorting of the genomic data. (C) Clustering the dataset’s samples and features using various algorithms and distance functions is available through the *Clustering* panel. Resulting clustering solutions are aggregated for future review and filtering. (D) The *Analysis* panel provides access to several visualization and exploratory tools like PCA, t-SNE, survival analysis, biomarker discovery, GO enrichment and automatic classifier generation. (E) In the *Dataset Collection* panel, several genomic matrices can be assembled into a multi-omic dataset collection, and then analyzed together.

## Results

We now describe PROMO’s main features, organized by analysis steps. The dataset used was TCGA’s breast cancer gene expression profiles (1218 samples downloaded from UCSC’s XENA website on May 2018). It is also available on the datasets page of PROMO’s website.

### Data import and preprocessing

In all analysis types, the first steps are to import the required data from local files into PROMO, and prepare it for the anlaysis. PROMO enables the integration of data of different types and from multiple sources by importing genomic matrices, sample labels and sample or gene partition files. Genomic matrices accompanied by complementary phenotypic information (clinical labels) can be loaded in the following formats: tabular text files, UCSC’s XENA[18][19] file formats (available for many public datasets including all TCGA’s data), and PROMO’s DSC files. The latter are precompiled multi-omic datasets available at PROMO’s dataset download page for selected TCGA cohorts. PROMO also allows separate loading of additional clinical labels and sample partition files to be used in the subtype discovery workflow.

After import, the loaded dataset can be ‘cleaned’ by filtering out samples based on clinical label values, and also by removing certain features (e.g., removing low variability genes or keeping only specific genes). Additional available common preprocessing steps include flooring, ceiling, and row normalization.

### Data exploration and visualization

Once a genomic matrix is loaded to PROMO, its properties can be explored with respect to any selected clinical label (Figure 3A). The samples (columns) in the matrix can be reordered based on any clinical label or by their mean expression. Basic dataset properties like value distribution (3B), clinical label distribution (3C), and sample variation (3D) can be studied and displayed graphically in various ways including PCA [20][21] and t-SNE [22]. For ease of interpretation, all displays consistently use the same colors to represent the various sample subgroups.

**Figure 3:**
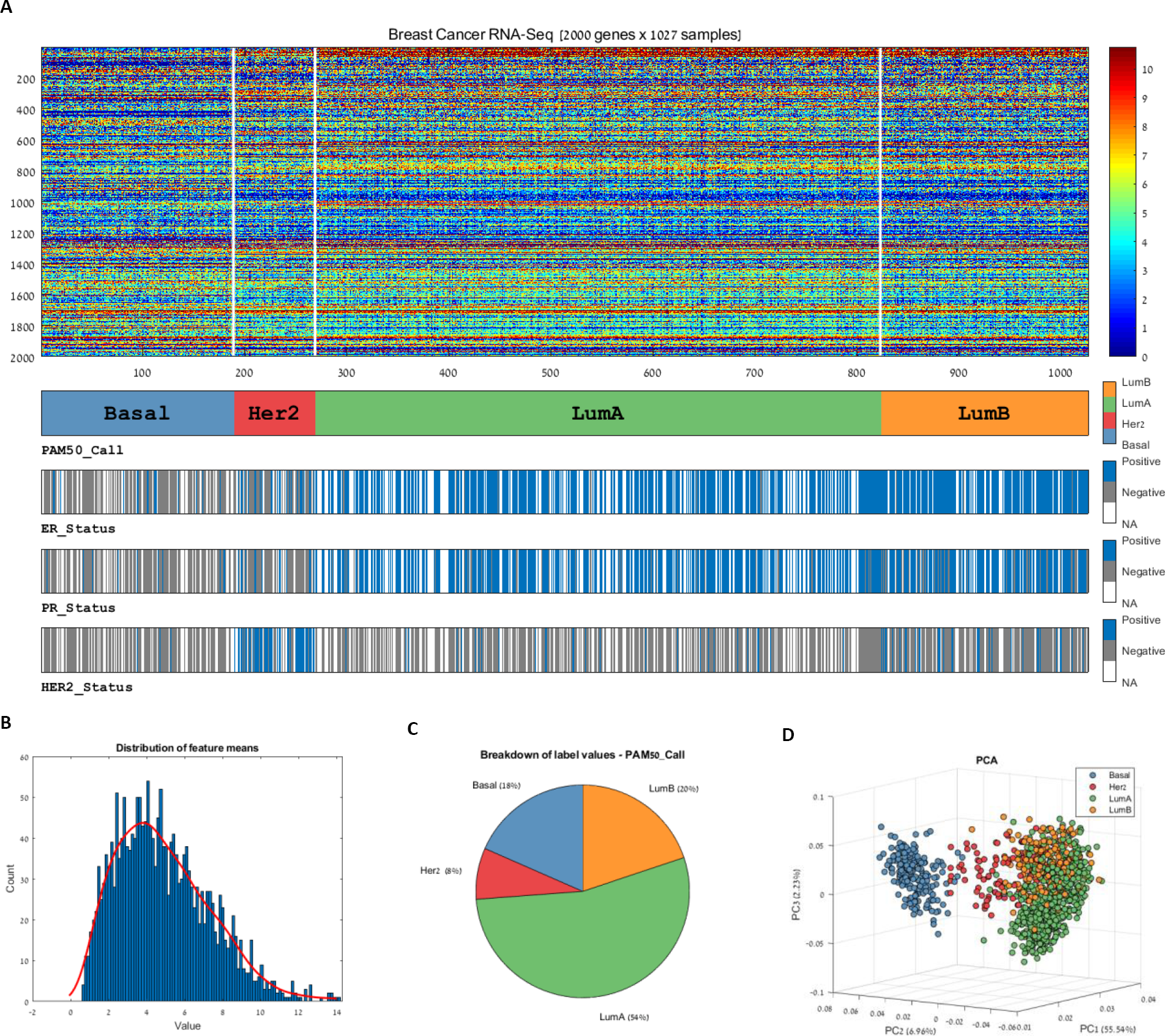
Visualization of multi-label genomic data. PROMO provides a variety of methods for visualizing a genomic dataset together with its associated clinical information. (A) A multi-label expression matrix plot. The plot is composed of a heat-map representation of the genomic matrix and several label bars beneath it showing different clinical labels that the user interactively selected. The colors in each label bar show the label value of each sample according to the legend on the right. The label appears below the lower left corner of the bar. Here, breast cancer patient profiles were grouped according to their PAM50 category (shown in the top label bar). By observing the distribution of values in other bars, relations between the groups and the labels can be observed. For example, the ER, PR and HER2 status of most samples in the ‘Basal’ group are negative, while the HER2 status of most ‘HER2’ group is positive. (B) Data distribution and (C) Clinical label distribution can be explored and visualized separately, or in combination using plots such as (D) PCA and others. These figures show that the Basal tumor samples are mainly characterized by Negative ER, PR and HER2 labels (A) and markedly differ from all other subtypes in their gene expression pattern (D), in accordance with the literature [14].

### Clustering and enrichment analyses

A major effort in promoting precision medicine is to identify disjoint groups of similar patients and characterize each group using its distinct genomic profile, survival data and clinical information. To reveal the similarities among patients, clustering is often performed on both samples and features [23]. Clustering the samples can reveal patient groups corresponding to disease subtypes [24], while clustering the features reveals groups of co-regulated genes [25]. PROMO provides various clustering algorithms such as K-means [26], hierarchical clustering [27], and Click [28] (PROMO’s clustering panel is shown in Figure S1). To explore the resulting clusters, the reordered matrix can be visualized in comparison to multiple sample labels (Fig 4A).

**Figure 4:**
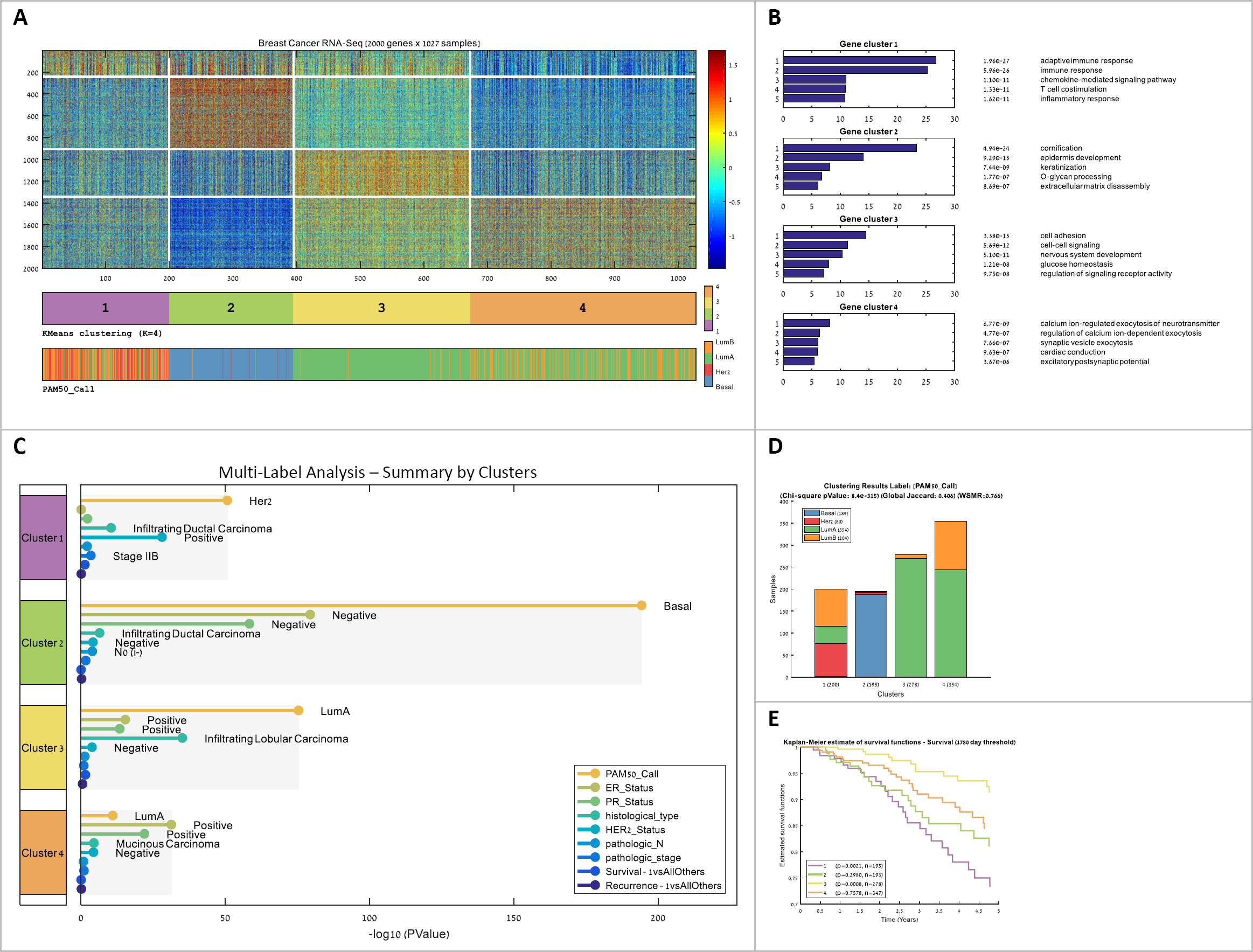
Identification and characterization of cancer subtypes. Unsupervised analysis followed by enrichment analysis is performed on both samples and features for identifying clinically significant samples groups, and for biologically characterizing them based on the functions of co-expressed gene groups. **(A)** The RNA-Seq expression matrix of TCGA’s breast cancer cohort after clustering both samples (columns) and genes (rows) into four clusters using the K-means algorithm. Clustering is based on the top 2000 variable genes. White lines separate clusters in each dimension. The bars below the matrix show selected sample labels (here: the clustering and PAM50). Matrix and bars were created using PROMO’s multi-label matrix drawing. **(B)** Gene clusters were characterized using PROMO’s gene ontology enrichment tool. The figure shows the five most significant GO terms for every gene cluster. **(C-E)** Sample clusters were characterized using the sample clinical labels: (C) PROMO’s multi-label analysis tool automatically tests the clinical labels of different types (numeric, ordinal, categorical or survival) for enrichment on the sample clusters. The various (D) Sample clusters can also be characterized for a single label by showing its value distribution in each cluster and by calculating enrichment. (E) Survival functions for each cluster. The p-values are the significance of separation of each cluster from the rest using the log-rank test.

After the genes have been clustered, the built-in Gene Ontology tool can help interpret the biological meaning of gene clusters using enrichment analysis (Fig 4B) [29]. Likewise, the clinical labels on the samples can be used to statistically characterize each sample cluster. A comprehensive analysis can be applied to each sample cluster using all clinical labels available for the cohort (numeric, ordinal, categorical or survival labels). The result is a characterization of each cluster together with FDR corrected p-values [30][31] in a unified report (Fig 4C). Enrichment tests for the sample clusters can also be performed using any selected single clinical label (Fig 4D). Finally, survival analysis performed on the sample clusters can test their prognostic value using Kaplan-Meier plots [32] and log-rank (Mantel–Haenszel) test [33](Fig 4E). Taken together, PROMO’s clustering and automatic multi-label enrichment analysis can quickly partition both samples and features into distinct groups and assess their biological meaning using the clinical labels.

### Identification of distinguishing genes and features (Biomarker discovery)

Having obtained patient subgroups of interest, either by sample clustering or using a predefined sample label, we may wish to identify distinguishing genes and features that differ significantly among sample groups. Such differentially expressed genes can shed light on the biological difference between sample clusters, and act as biomarkers for classifying a new sample to a sample class.

After selecting the label and the groups that will be compared, PROMO enables the application of various statistical tests for identifying genes that are differentially expressed among the groups. The p-values obtained by the tests can be used for gene sorting, filtering and for clustering the genes into up-regulated and down-regulated groups. PROMO’s Gene Ontology enrichment analysis can be executed on the resulting gene groups for characterizing the function of up-regulated and down-regulated genes. FDR correction and fold-change based filtering are also supported. PROMO’s biomarker discovery panel and an example of its output are shown in Figure S2.

For detecting survival biomarkers, PROMO can rank all genes by their association to survival, based on Cox regression analysis [34]. In addition, the user can use the expression levels of selected genes to generate a new sample label (for example HER2_Low and HER2_High). Kaplan-Meier plots can then be used to estimate the significance of survival differences between sample groups defined by the new label.

Lastly, PROMO can help in finding genes that are functionally related to a given gene of interest by ranking all genes based on their correlation to it. Altogether, the various techniques described here and implemented in PROMO can quickly identify genes that take part in the biological differences between sample groups and may serve as biomarkers for the selected label.

### Automatic generation of a simple molecular classifier

After having partitioned the dataset samples, characterized the sample groups and their genes, and established the clinical relevance of the groups, PROMO can build an algorithm to classify a new sample into one of the groups. Such a classifier, especially if based on a small number of genes (rather than the thousands used to identify the subgroups) can serve as a significant step towards translating the analysis results into a diagnostic biomarker for clinical use.

Of the many possible classifier types, decision trees have the advantages of being easy to understand, highly interpretable biologically and easily visualized [35]. Furthermore, they allow for controlling the tradeoff between accuracy and simplicity. For predicting any selected sample label, PROMO can generate a simple decision tree with a single click (Fig 5). The generated decision tree can be visualized graphically, specified textually and saved to a Matlab file as a function. Automatic cross-validation and parameter optimization make it easy for the user to come up with a simple decision tree that may be in future subtype classification kits. It is also possible to generate a large number of random trees and rank the genes by the frequency of their appearance in the trees, thus identifying informative features for subtype classification.

**Figure 5:**
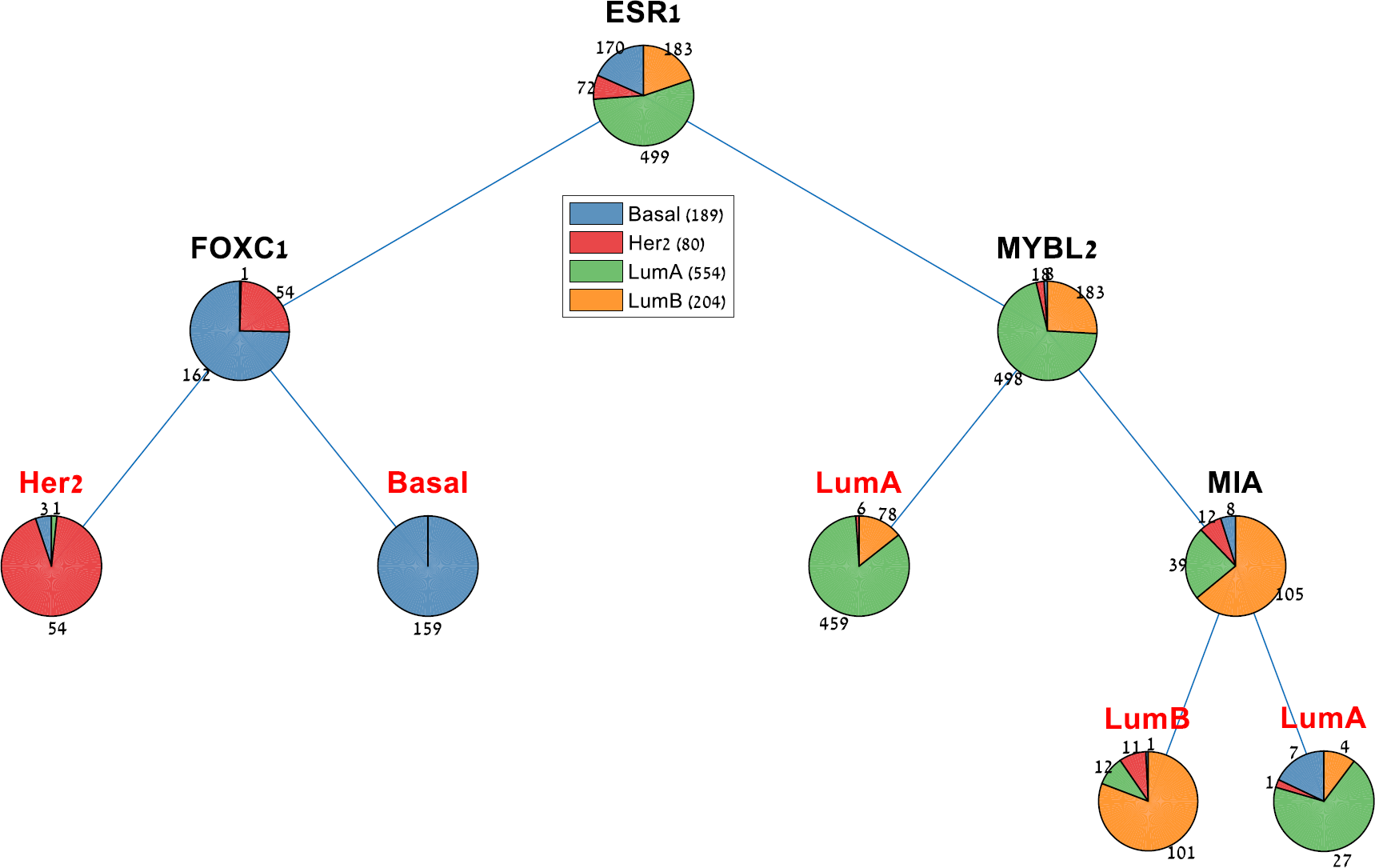
Automatically generated decision tree for classifying breast tumors into the four PAM50 classes. PROMO can generate a cross-validated decision tree for any selected sample label using the currently loaded matrix as training data. In this figure, a four-gene molecular classifier for breast cancer subtypes is presented, showing 7.77% loss on the training data, and 15% averaged loss on 10-fold cross-validation.

### Integrative multi-omic analysis

In multi-omic datasets, each sample is characterized by several omic profiles (e.g., gene expression, methylation, copy number). Integrative analysis of multi-omic cancer datasets has a potential of revealing biological regulatory patterns that are missed in single omic analysis, and tools for performing such analyses are currently in great demand [36][37].

PROMO provides several features for handling and analyzing multi-omic datasets. The profiles composing a multi-omic dataset can be imported from repositories into a ‘Multi-Omic Dataset Collection’ in PROMO (Figure 2E). The user can navigate between the matrices, edit them independently, and select a subset of the datasets for downstream integrative analysis. Precompiled dataset collections for several TCGA cancer type cohorts are available on PROMO’s download page.

After setting up a multi-omic collection, the “inter-omic correlation identification” feature helps to detect correlations between features in two selected omics. This feature allows the identification of correlations between features from different biological levels. For instance, anti-correlation between mRNA expression and DNA methylation levels can pinpoint biological regulation.

The “Multi-omic clustering” feature can be used to cluster the dataset samples based on several omic matrices simultaneously. To this end, PROMO provides implementations of the multi-omic algorithms SNF [38], NEMO [39] and of Consensus Clustering [40] modified for multi-omic data. Figure S4 demonstrates the application of a multi-omic clustering algorithm on three different omics of the TCGA’s breast cancer cohort.

## Discussion

Recent cancer projects such as TCGA [41], GDC [42], and ICGC [43] provide the research community with a wealth of omic profiles and extensive clinical information on cancer patients [44]. Analysis of the data is challenging and requires advanced bioinformatics, statistical and programming skills. A thorough analysis of these datasets - and larger ones expected in the future - by many researchers, is crucial for improving cancer diagnosis and treatment.

PROMO aims to fill in a gap in available analysis tools for such large genomic and clinical cancer datasets. It is an interactive tool that is freely available and supports a rich collection of analysis methods and facilitates useful workflows for data exploration and visualization, cancer subtype identification, biomarker discovery and integrative multi-omic analysis. (See Table 2 for a list of the key features). PROMO’s support for large sample size in addition to features like survival analysis and interrogation of the clinical data on sample clusters make it especially suitable for analyzing modern cancer datasets. While many of PROMO’s features are also available in other tools (Table 3), PROMO is unique in its comprehensiveness, support for large sample dimension and the spectrum of tools it provides.

**Table 2:** PROMO’s key features

| CATEGORY | KEY FEATURES |
| --- | --- |
| DATA IMPORT | ▪ Importing genomic data from tabular CSV files<br>▪ Importing UCSC's XENA genome matrix and phenotype files<br>▪ Adding clinical labels from file |
| PREPROCESSING | ▪ Flooring, ceiling and row normalization<br>▪ Filtering of samples by clinical labels<br>▪ Filter features by range, variance, gene symbols or by an external list |
| DATA EXPLORATION AND VISUALIZATION | ▪ PCA, t-SNE<br>▪ Data distribution plots<br>▪ Survival Analysis (Kaplan Meier, Log rank)<br>▪ Multi-label expression matrix figures |
| SORTING | ▪ Sorting samples and features based on genomic data<br>▪ Sorting samples based on clinical labels |
| CLUSTERING | ▪ Clustering both samples and features using K-means [26], hierarchical clustering [27], and Click [28]<br>▪ Browsing clustering history and zooming into specific clusters |
| SAMPLE CLUSTER ANALYSIS | ▪ Automated multi-label enrichment test for detecting enrichment of clinical labels |
| FEATURE CLUSTER ANALYSIS | ▪ Gene ontology enrichment analysis |
| BIOMARKER DISCOVERY | ▪ Applying statistical tests for detecting differentially expressed genes/features<br>▪ Filter results by FDR corrected p-value and fold change<br>▪ Rank genes based on survival prediction (COX regression) |
| CLASSIFIER GENERATION | ▪ Automatic generation of decision tree classifiers for selected sample labels |
| INTEGRATIVE MULTI-OMIC ANALYSIS | ▪ Assembly of dataset collection<br>▪ Multi-omic clustering using SNF [38], NEMO [39] or Consensus Clustering [40]<br>▪ Inter-omic correlation identification |

**Table 3:** Comparison of the main functions provided by PROMO and by other tools

| **Function** | **PROMO** | **Expander**<br>[45] | **XENA**<br>[19] | **Perseus**<br>[46] | **KnowEng**<br>[47] | **O-Miner**<br>[48] |
| --- | --- | --- | --- | --- | --- | --- |
| *Precompiled datasets* | V | X | V | X | V | V |
| *Preprocessing* | V | V | X | V | X | V |
| *Data Visualization* | V | V | V | V | V | V |
| *Sample clustering* | V | V | X | V | V | V |
| *Feature clustering* | V | V | V | V | X | V |
| *Sample clusters enrichment tests (clinical data)* | V | X | V | X | V | X |
| *Feature clusters enrichment tests* | V | V | X | V | V | V |
| *Survival analysis* | V | X | V | X | V | V |
| *Biomarker discovery* | V | V | X | V | X | V |
| *Automatic decision tree generation* | V | X | X | X | X | X |
| *Inter-omic correlation identification* | V | X | X | X | V | X |
| *Integrative multi-omic sample clustering* | V | X | X | X | X | X |

Our vision for PROMO is that it will be used as a one-stop shop for mining clinically important insights from genomic datasets, quickly and without any need for programming skills. It accelerates the analysis process and makes it more accessible for non-computational cancer researchers. Within a single short session, the user can import a cancer dataset of interest, preprocess it, cluster its samples and features, test the sample clusters for significance using survival analysis and enrichment tests on the clinical labels, test the feature clusters for GO enrichment, identify subtype distinguishing features (biomarkers) using various statistical tests and export the results using various reports and figures. The simple classification capabilities in PROMO can automatically produce a decision tree classifier for any selected label, and thus act as a basis for a subtype diagnosis.

We intend to continue developing PROMO by adding features and supporting the tool’s users. We hope that PROMO’s comprehensiveness and ease of use will help cancer researchers make the best use of the accumulating cancer datasets to fulfill the promises of precision medicine.

## Conclusions

PROMO is a powerful, user-friendly, stand-alone, publicly available tool for exploration, analysis, and interpretation of genomic cancer data together with clinical information.

## Availability and requirements

Project name: PROMO (Profiler of Multi-Omics data)

Project home page: http://acgt.cs.tau.ac.il/promo/

Operating system: Windows

Programming language: Matlab

Required runtime library: Matlab R2019a (9.6)

## List of abbreviations

BRCA: Breast Cancer
DSC: Dataset Collection
FDR: False Discovery Rate
GO: Gene Ontology
PCA: Principal Component Analysis
t-SNE: t-distributed Stochastic Neighbor Embedding
PROMO: Profiler of Multi-Omics data
TCGA: The Cancer Genome Atlas

## Declarations

The results published here are based upon data generated by The Cancer Genome Atlas managed by the NCI and NHGRI. Information about TCGA can be found at http://cancergenome.nih.gov.

## Ethics approval and consent to participate

Not applicable

## Consent for publication

Not applicable

## Availability of data and material

Software and data available at http://acgt.cs.tau.ac.il/promo/

## Authors’ contributions

DN designed and wrote the code. NS contributed to the design and implementation. IL contributed to the implementation. RS supervised the study. DN and RS wrote the manuscript.

## Acknowledgements

Not applicable

## Supplementary Material

**Figure S1:**
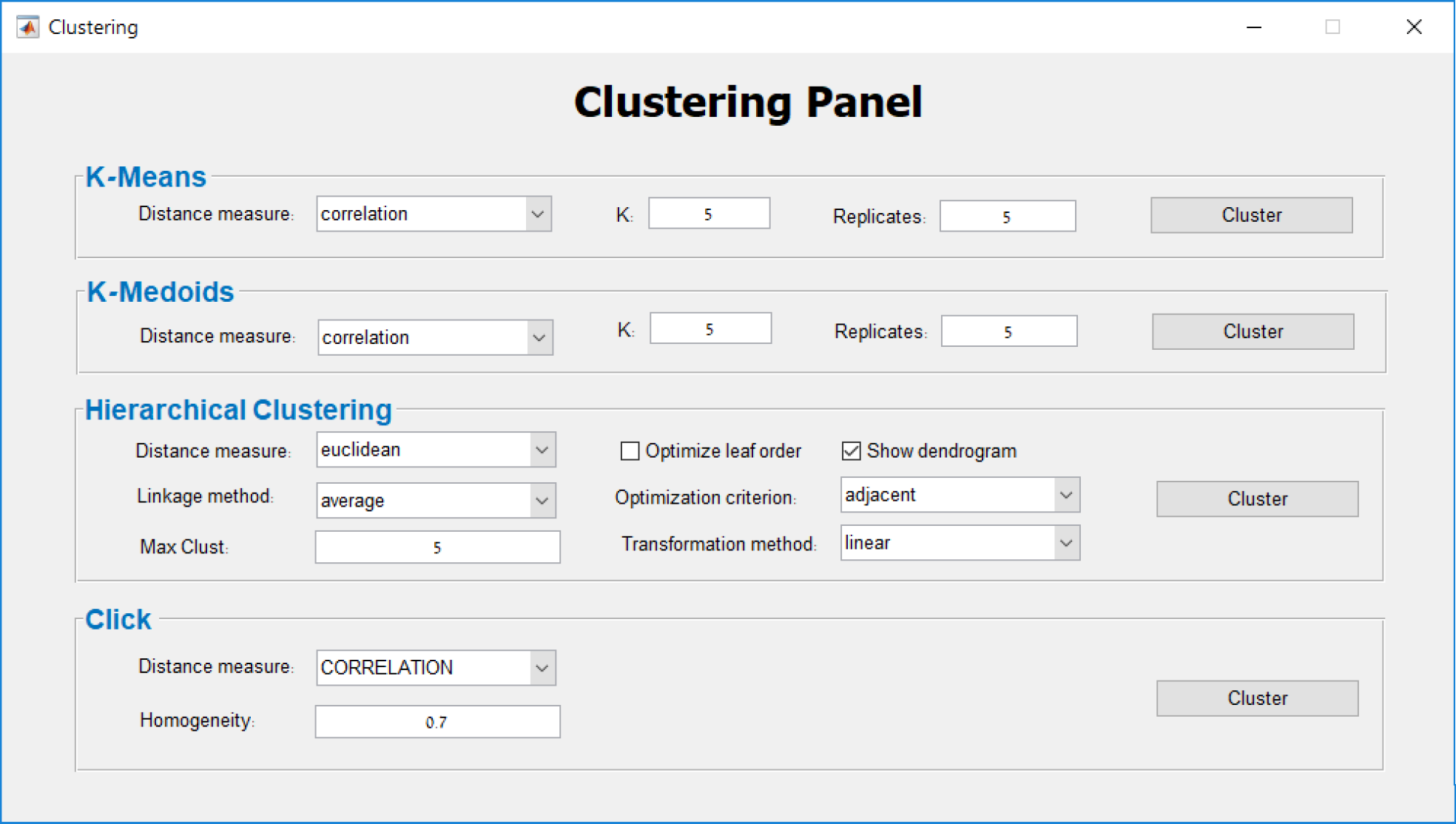
Clustering Panel. The clustering panel allows the selection of a clustering algorithm and its relevant parameters. Clustering can be applied both on samples and on genes. The resulting clusters are added as a new sample label and can be explored on PROMO’s main screen with respect to any other clinical label (See Figure 3).

**Figure S2:**
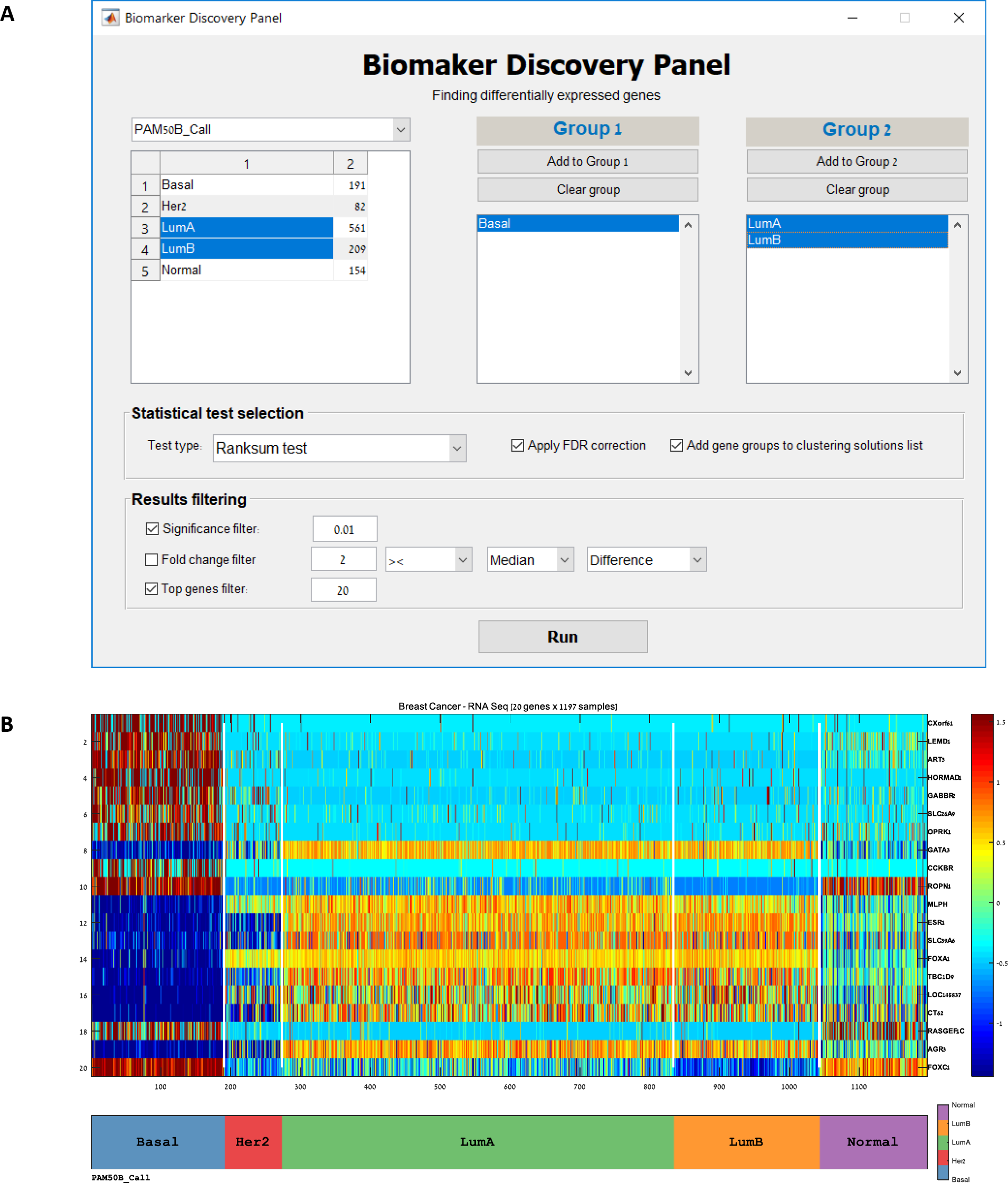
Biomarker Discovery. (A) This panel is used for identifying genes that are differentially expressed between sample groups defined by any sample label. Statistical tests vary by the label types, and include t-test, Ranksum test, ANOVA and Kruskal-Wallis. After optional filtering, the resulting list of genes is saved to a file sorted by p-value. Here two groups were defined, according to the PAM50 label. One group corresponds to the basal and the other to the LumA and Lum B categories. See Table S1 for the resulting set of differentially expressed genes. (B) The feature patterns of the identified genes are presented on PROMO’s main screen together with any selected sample labels. Here we see the expression levels of the 20 genes that were identified by the test in A, after row normalization).

**Table S1:** List of differentially expressed genes. The 20 genes with the most significant differential expression between the groups defined in Figure S2A are shown. Genes are sorted by their Ranksum test p-values. Genes with positive fold change are over-expressed on the Basal samples compared with the Luminal samples. Here, for instance, we see that the Estrogen Receptor gene (ESR1) is ranked 12^th^ and exhibits a significant under-expression on the Basal tumors samples (the Triple-Negative subtype) compared to the Luminal tumor samples.

| *id* | *Gene Symbol* | *p-value (Test:*<br>*Ranksum test on*<br>*PAM50B_Call*<br>*[Basal](n=191) vs.*<br>*[LumA,LumB](n=770))* | *Fold*<br>*Change* | *p-value*<br>*Rank* |
| --- | --- | --- | --- | --- |
| 1 | CXorf61 | 2.33E-123 | 4.5604 | 1 |
| 2 | LEMD1 | 1.72E-122 | 3.1005 | 2 |
| 3 | ART3 | 3.17E-118 | 5.294 | 3 |
| 4 | HORMAD1 | 1.75E-113 | 5.8894 | 4 |
| 5 | GABBR2 | 7.04E-111 | 4.2383 | 5 |
| 6 | SLC26A9 | 4.21E-101 | 2.4335 | 6 |
| 7 | OPRK1 | 2.22E-99 | 2.5534 | 7 |
| 8 | GATA3 | 3.27E-99 | -4.02715 | 8 |
| 9 | CCKBR | 5.86E-99 | 2.1373 | 9 |
| 10 | ROPN1 | 8.44E-99 | 6.1879 | 10 |
| 11 | MLPH | 5.43E-98 | -5.1038 | 11 |
| 12 | ESR1 | 7.07E-98 | -7.07625 | 12 |
| 13 | SLC39A6 | 9.68E-98 | -2.7197 | 13 |
| 14 | FOXA1 | 3.21E-97 | -6.7128 | 14 |
| 15 | TBC1D9 | 6.71E-97 | -4.06355 | 15 |
| 16 | LOC145837 | 2.99E-96 | -4.18775 | 16 |
| 17 | CT62 | 3.12E-96 | -3.65845 | 17 |
| 18 | RASGEF1C | 5.50E-96 | 2.1274 | 18 |
| 19 | AGR3 | 1.12E-95 | -9.2111 | 19 |
| 20 | FOXC1 | 1.63E-95 | 4.26835 | 20 |

**Figure S3:**
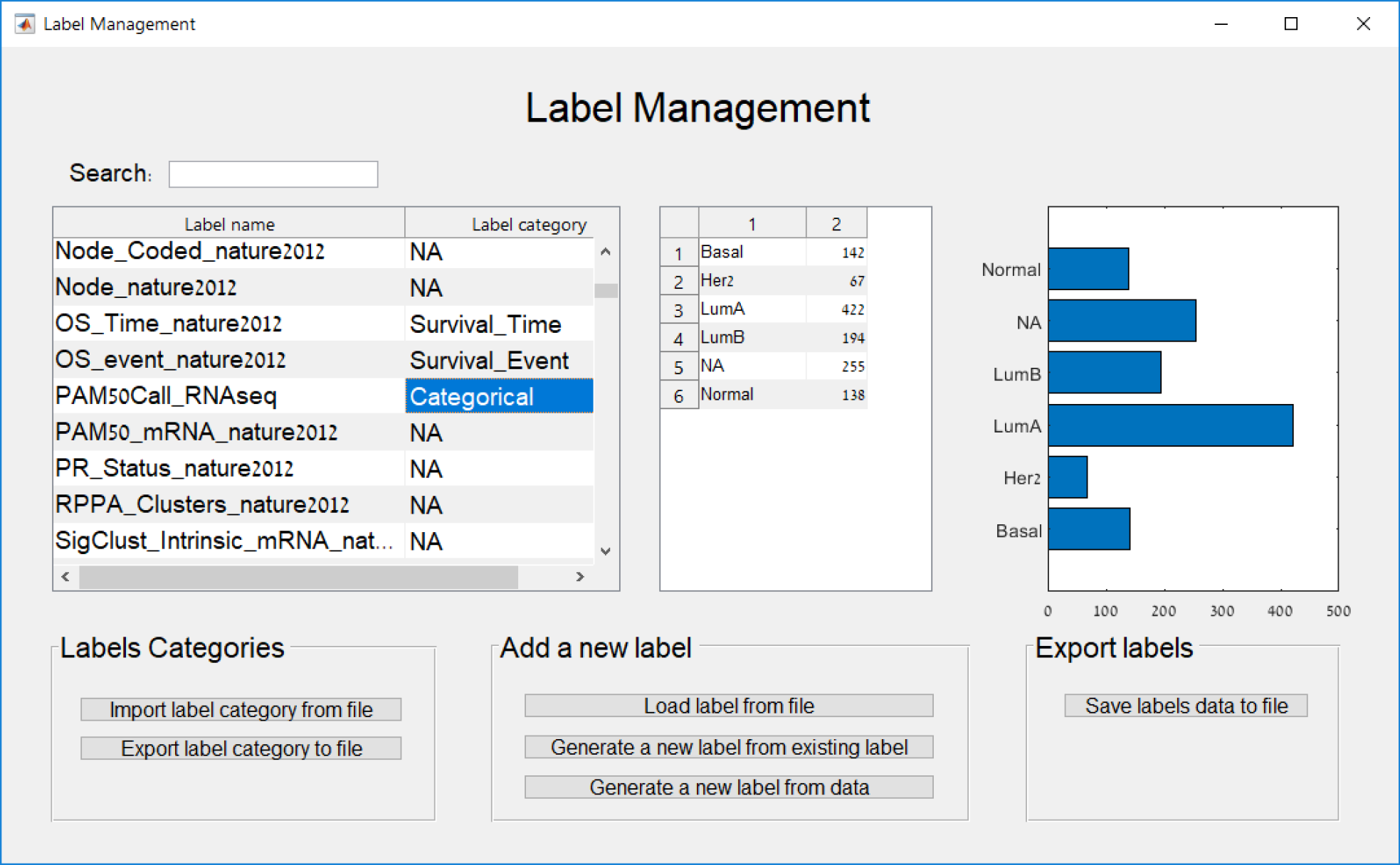
Label Management Panel. This panel allows the management of sample labels, including removing, renaming and viewing the distribution of values of a label. Labels can be assigned to category types, and those types determine the statistical test that can be used for calculating their enrichment on sample clusters. Both labels and their categories can be loaded and saved to files. New labels can be generated from existing labels (by uniting label values for instance), or from genomic data (e.g., translating expression values of selected gene to LOW/HIGH labels). Lastly, the distribution of values for the selected label is displayed as an histogram on the right.

**Figure S4:**
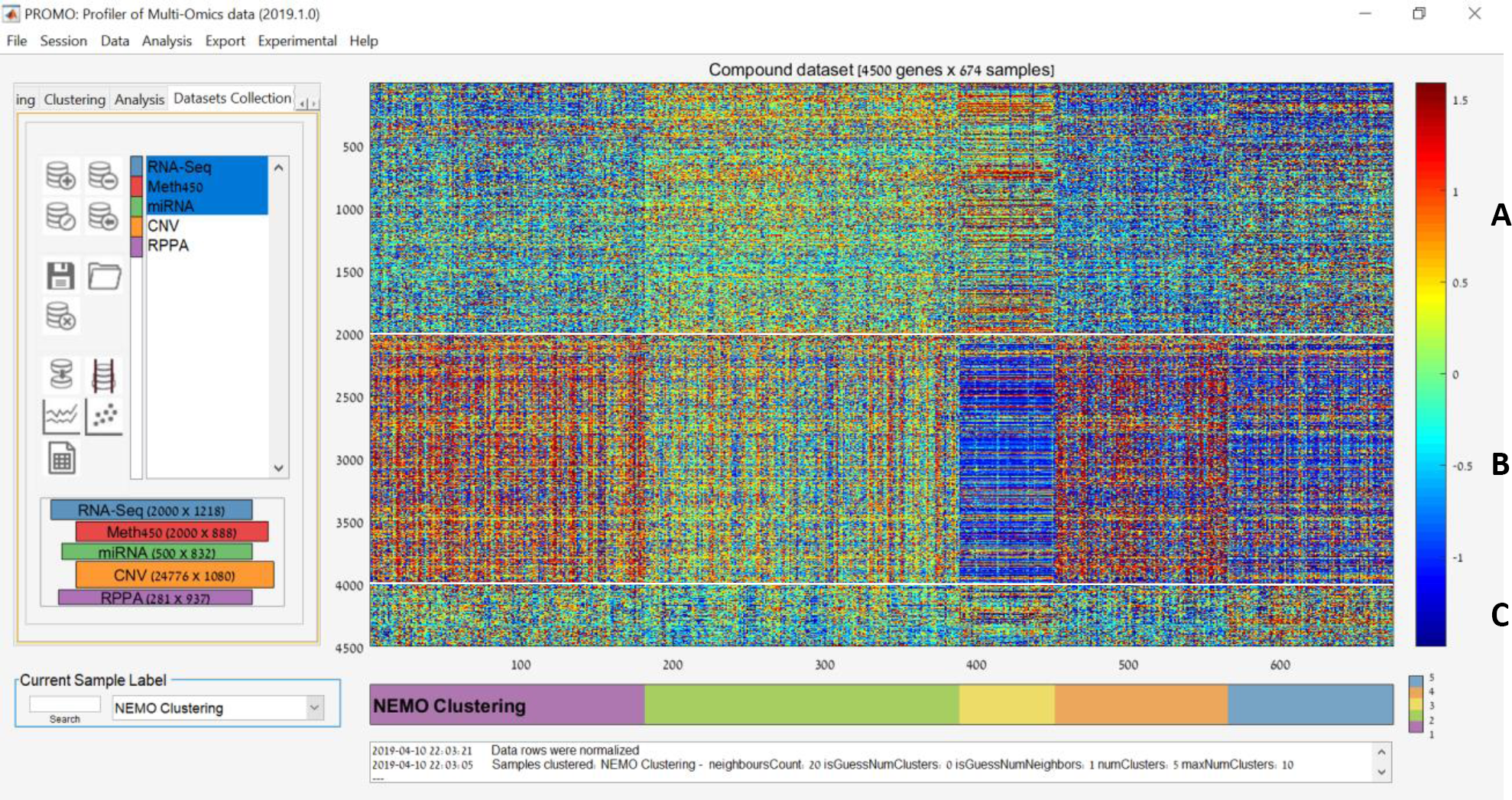
Multi-omic sample clustering. Screenshot of PROMO’s main screen after applying multi-omic clustering on 674 breast tumor samples from TCGA. The ‘Dataset Collection’ panel on the left was used to select the three omics to be used in the clustering. Here features from three different omics were used: (A) RNA-Seq (2000 features), (B) DNA methylation arrays (2000 features) and (C) miRNA arrays (500 features). Algorithm NEMO [39] was applied on the subset of samples appearing in the three omics into 5 groups, shown on the label bar below the matrix. The genomic matrix displays concatenation of the 4500 features included in the analysis after row normalization, with samples grouped by cluster. The 1^st^ and 4^th^ clusters from the left have high methylation signals, while the second and third have higher gene expression signals. Clustering of tumor samples using a multi-omic algorithms integrates data from different biological levels and thus has the potential of revealing biological regulatory patterns that are missed in single omic analysis.

